# Pharmacological BET Protein Inhibition Reverts Niemann-Pick Type C Disease-Associated Changes in Human iPSC-Derived Cortical Neurons

**DOI:** 10.64898/2026.08.31.747771

**Authors:** F. Benigni, M. Parente, C. Marioli, R. Borghi, N. Cicolani, A. Barthelemy, S. Petrini, M. Tartaglia, FW. Pfrieger, V. Pallottini, C. Compagnucci

**Affiliations:** Molecular Genetics and Functional Genomics, Ospedale Pediatrico Bambino Gesù, IRCCS, 00146 Rome, Italy; Department of Science, Section Biomedical Science and Technology, University Roma Tre, Viale Marconi 446, 00146 Rome, Italy; Microscopy Core Facility, Research Center, Bambino Gesù Children’s Hospital, IRCCS, 00146 Rome, Italy; Centre National de la Recherche Scientifique, Université de Strasbourg, Institut des Neurosciences Cellulaires et Intégratives; 8 allée du Général Rouvillois, 67000 Strasbourg, France; LIFE Institute for Research and Health Care - Santa Lucia IRCCS, Via del Fosso Fiorano 64, 00143 Rome, Italy

**Keywords:** Niemann-Pick type C1, lysosomal storage disorder, cholesterol trafficking, iPSC-derived neurons, JQ1, epigenetic regulation, neurodegeneration, bromodomain proteins, histone acetylation

## Abstract

Niemann-Pick type C1 (NPC1) disease is a fatal lysosomal disorder caused by impaired intracellular cholesterol trafficking, leading to progressive neurodegeneration and premature death. Despite advances in disease modelling, therapeutic development remains limited, partly due to the lack of human neuronal systems that accurately recapitulate disease-relevant phenotypes.

Here, we generated a patient-specific neuronal model by differentiating induced pluripotent stem cells (iPSCs) homozygous for the most common *NPC1* c.3182T>C (p.I1061T) variant into cortical neurons. These cells exhibited hallmarks of neuronal pathology associated with NPC1 disease. Using this experimental model, we explored the effects of pharmacological inhibition of bromodomain and extraterminal (BET) proteins. Treatment with the BET inhibitor JQ1 attenuated disease-associated endophenotypes, improving neuronal viability and modulating intracellular cholesterol accumulation. Overall, our findings identify BET inhibition as a novel strategy to ameliorate NPC1-associated neuronal pathology, uncover a previously underexplored link between epigenetic regulation and cholesterol homeostasis, and highlight the use of iPSC-derived neurons as a powerful platform for therapeutic discovery.

## Introduction

Niemann-Pick disease type C (NPCD) is a rare autosomal recessive neurovisceral lysosomal disorder characterized by progressive neurodegeneration, hepatosplenomegaly, and reduced lifespan [1–5,7,43]. It is caused by biallelic pathogenic variants in either the *NPC intracellular cholesterol transporter 1 (NPC1*; OMIM #257220) or *NPC intracellular cholesterol transporter 2 (NPC2*; OMIM #607625) genes, with approximately 95% of affected individuals carrying mutations in *NPC1* [1,5–10,92]. *NPC1* encodes a transmembrane protein localized to the limiting membrane of late endosomes and lysosomes, whereas *NPC2* encodes a soluble lysosomal cholesterol-binding protein [11–15,21,32]. Together, these proteins mediate the exit of unesterified cholesterol from the endosomal-lysosomal system, enabling its redistribution to other cellular compartments, including the endoplasmic reticulum and plasma membrane [12,13,16,17,19–21,39]. Loss of NPC1/NPC2 function disrupts this pathway, resulting in lysosomal accumulation of cholesterol and other lipids [19,27,40,41,62]. This primary defect triggers widespread cellular dysfunction, including impaired autophagy, mitochondrial abnormalities, altered lysosomal homeostasis, and compromised neuronal survival [25,28,29,30,34,36,44,46,47,50]. Neurons are particularly vulnerable due to their postmitotic state, high metabolic demand, and dependence on efficient intracellular trafficking. As a result of impaired NPC1/NPC2 function, affected individuals show progressive neurodegeneration characterized by axonal transport defects and neuroinflammation [29,30,33–36,40,45,50–57,61].

Current therapeutic approaches in NPCD is mainly directed to target downstream metabolic consequences rather than the primary genetic defect [5,7,58–65]. Despite significant research efforts, current treatment options for NPCD remain limited to palliative care and a few disease-modifying agents, including N-butyl-deoxynojirimycin (OGT918, Miglustat, Zavesca) [66], arimoclomol (Miplyffa) in combination with Miglustat [67], and N-acetyl-L-leucine (Levacetylleucine) [68,69].

Our previous studies indicated bromodomain and extra-terminal (BET) proteins as potential drug target for NPCD [91,92]. BET proteins are histone acetylation readers that regulate transcriptional programs involved in inflammation, metabolism, and cellular stress responses [79–88,90], which are processes relevant to NPCD pathophysiology [78,89,91,92]. Pharmacological BET inhibition has shown beneficial effects in diverse disease models, although mechanisms are context-dependent and not fully understood [81,87,88,90,91]. In non-neuronal systems, BET proteins have also been implicated in lipid metabolism and lysosomal function [89,91], but their role in NPCD-associated neuronal dysfunction remains unclear.

Patient-derived induced pluripotent stem cells (iPSC) differentiated into cortical neurons were used as experimental model system. As previously shown, these neurons recapitulate key NPC phenotypes, including cholesterol accumulation, impaired cellular homeostasis, and dysregulated stress responses [70,71,73–76,98,99]. Although NPC is primarily considered a lipid trafficking disorder, emerging evidence suggests that transcriptional and epigenetic dysregulation may also contribute to disease progression, although these mechanisms remain incompletely defined [78,81,86–88,92].

To assess the effects of pharmacological BET inhibition in NPCD, we investigated the rescue of disease-associated endophenotypes in NPC1-deficient neurons, using JQ1, a well-characterized and widely used BET bromodomain inhibitor [79,83,88,90,92]. We generated cortical neurons from iPSCs derived from a patient carrying the common NPC1 p.I1061T amino acid substitution accounting for 20% of affected individuals [5,93,95,97], and the effects of BET inhibition on intracellular cholesterol accumulation, apoptotic susceptibility, and NPC1 protein expression was evaluated. The collected data documented an improved neuronal viability and reduced intracellular cholesterol accumulation, indicating a BET inhibition as a novel strategy to ameliorate NPC1-associated neuronal pathology.

## Results

### 1. Characterization of human iPSCs-carrying the I1061T variant of NPC1

An iPSC line from the Coriell Biobank generated from primary fibroblasts carrying the homozygous c.3182T>C change (p.I1061T) in *NPC1*, and one from fibroblasts of a healthy donor were used in the study. Both lines were characterized using established assays. Colonies of selected and expanded iPSCs resembled embryonic stem cells (ESCs) and exhibited a round shape with clearly delineated borders without noticeable differences between genotypes (Fig. 1A). Immunocytochemical staining with informative markers and the alkaline phosphatase (ALP) assay confirmed pluripotency of the colonies regardless of their genotype (Fig. 1B-D).

**Figure 1:**
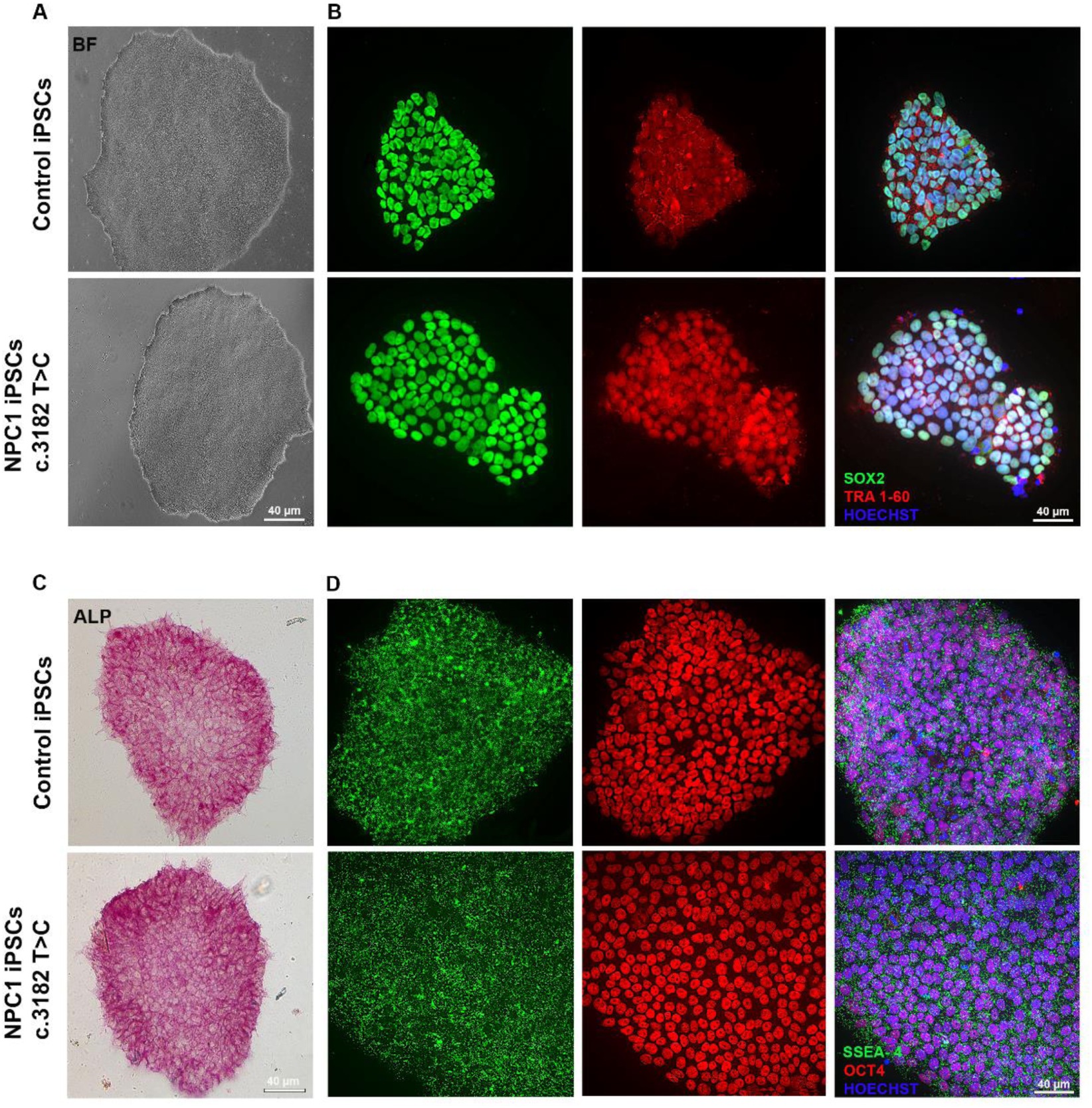
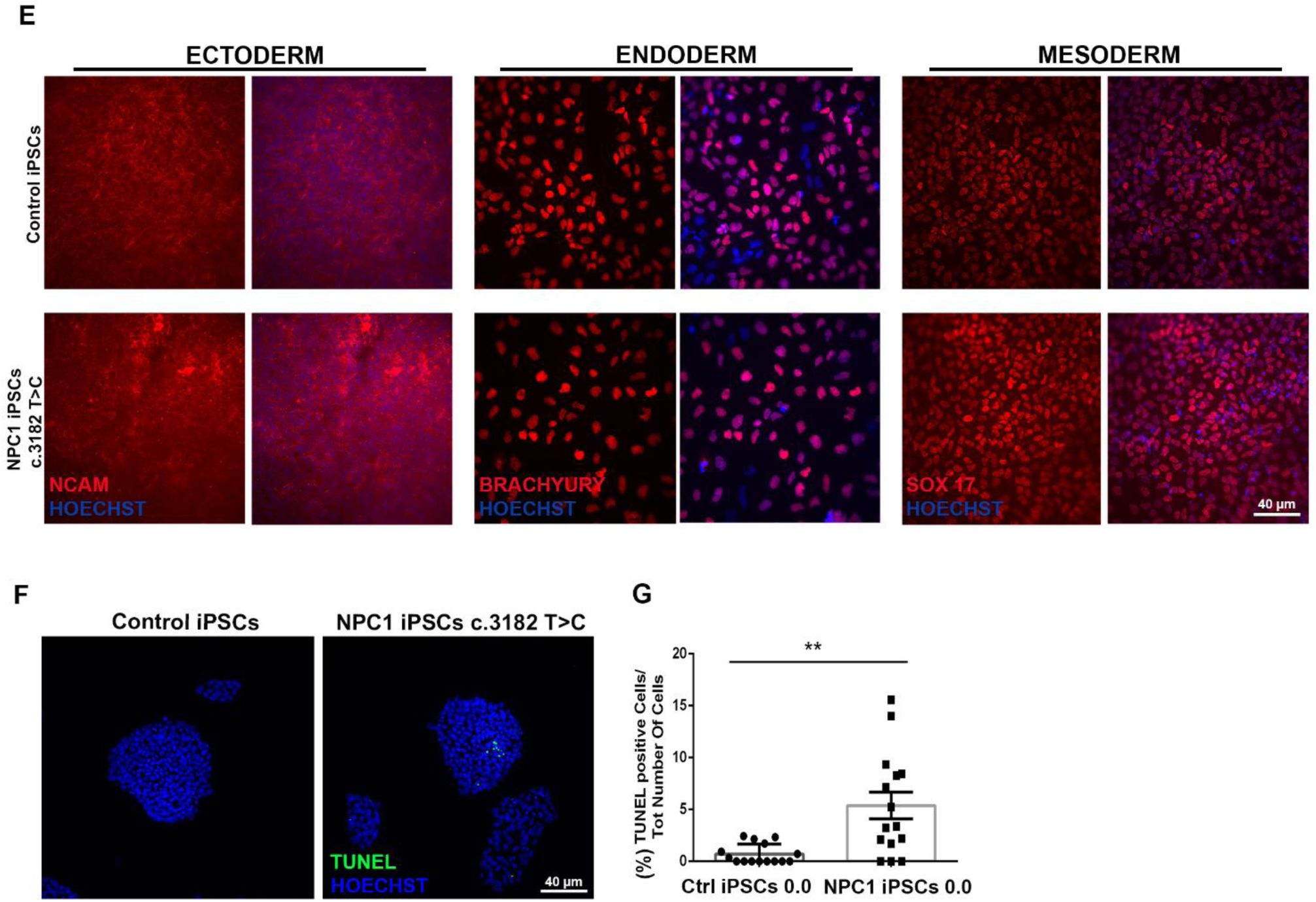
Characterization of iPSCs obtained from NPC1 cells. (**A**) Representative brightfield micrographs of control and NPC1-mutated iPSCs forming colonies with rounded shape. Scale bar: 40 μm. (**B, D**) Representative immunofluorescence micrographs of colonies following immunocytochemical staining of pluripotency markers SOX2 and SSEA4 (green), TRA1-60 and OCT4 (red). Nuclei were counterstained with Hoechst (blue). Scale bar: 40 μm. (**C**) Bright-field micrographs of iPSC colonies showing presence of alkaline phosphatase (ALP). Scale bar: 40 μm. (**E**) Fluorescence micrographs of iPSC colonies showing presence of established markers for germ layer lineages (ectoderm: NCAM; endoderm: SOX17; mesoderm: BRACHIURY, all red) following trilineage assay and immunocytochemical staining. Nuclei are counterstained with Hoechst (blue). Scale bar: 40 μm. (**F**) Confocal micrographs of iPSCs cultures following TUNEL assay (green) and nuclear staining with Hoechst (blue). Scale bar: 40 μm. (**G**) Bar graphs showing percentage of cells with fragmented DNA in Hoechst-positive cells in NPC1 and control iPSCs colonies (whiskers indicate SEM). Percentages are reported as mean of n = 3 biological replicates. Asterisks indicate statistically significant differences (**, p ≤ 0.01; unpaired, two-tailed Student’s t-test).

The ability of iPSCs to differentiate to three germ layers was tested by the trilineage differentiation assay. In both lines, immunocytochemical staining revealed the presence of germ layer markers SOX17 (endoderm), BRACHIURY (mesoderm and NCAM (ectoderm; Fig. 1E). TUNEL assay revealed increased apoptosis in patient-derived iPSCs compared to controls (Fig. 1F,G). Together, these data revealed that the *NPC1* variant does not affect pluripotency and germ cell layer differentiation of iPSC colonies but decrease viability.

### 2. NPCD neurons exhibit morphological features comparable to control neurons

As next step, iPSCs were differentiated into cortical neurons (CNs) following a previously established protocol [98] (Fig. 2A). Progress of neuronal differentiation was monitored at several time points. At 10 days post-differentiation (DPD), cells exhibited neuronal polarization and neurite extension, progressively developing into mature neurons forming dense networks of neurites by 30 DPD.

**Figure 2:**
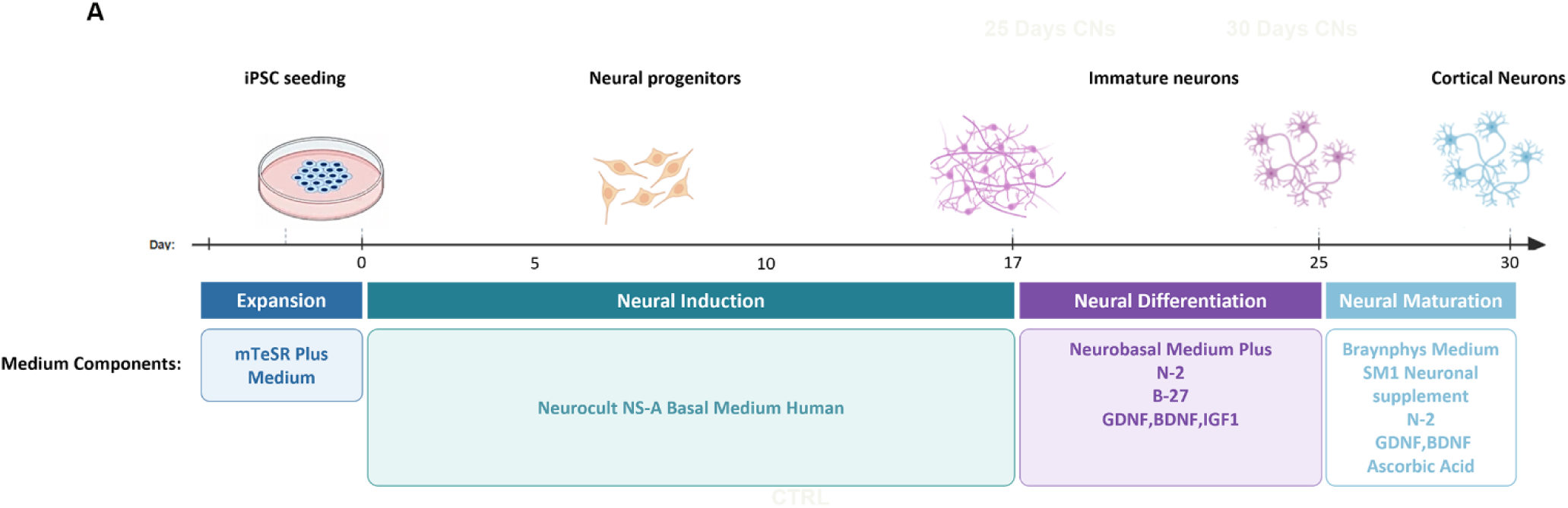
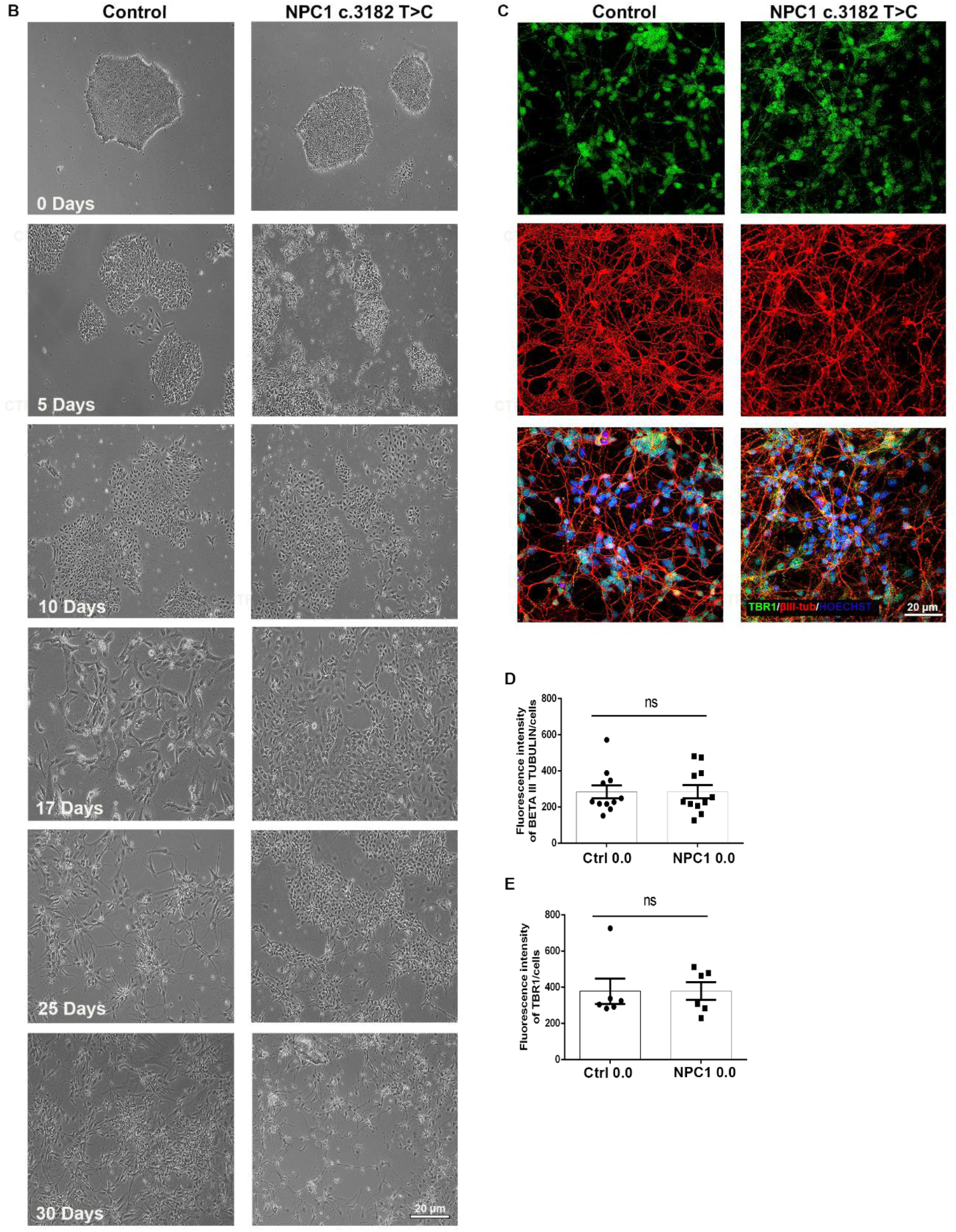
Differentiation of cortical neurons from control and NPC1 iPSCs. **(A)** Schematic representation of the iPSC-to-cortical neuron differentiation protocol, illustrating the sequential stages of iPSC expansion, neural induction, neuronal differentiation, and neuronal maturation, together with the culture media and supplements used at each stage. Generated using Biorender. **(B)** Cells were exposed to defined culture media and factors to promote neural progenitor formation (days 5-10) followed by maturation into cortical neurons (day 30). (**C**) Representative fluorescence micrographs showing presence of the cortical neuronal marker TBR1 (green) and the pan-neuronal marker βIII-tubulin (red) in neurons differentiated from NPC1 and control IPSCs. Nuclei were counterstained with Hoechst (blue). Scale bar: 20 μm. **(D**,**E)** Bar graph showing mean fluorescence intensity of indicated markers in somata of cortical neurons at 30 DPD. Whiskers indicate SEM. No statistical significant differences were observed between genotypes (n = 3 biological replicates; two-tailed Student’s t-test).

Notably, the morphology of neurons was comparable between control and NPCD cultures, indicating that the *NPC1* variant does not impair differentiation and structural integrity at the time points studied (Fig 2B). Immunocytochemical staining confirmed successful cortical specification, as demonstrated by the presence expression of T-box brain transcription factor 1 (TBR1), a marker of cortical neurons, and the pan neuronal marker βIII-tubulin (TUJ1) (Fig. 2C). Quantification (Fig. 2D-E) revealed no significant differences between control and patient-derived neurons with respect to neuronal marker expression supporting the generation of comparable neuronal populations suitable for subsequent analyses.

### 3. JQ1 does not alter morphology of control or NPC1 neurons

To evaluate the therapeutic potential of JQ1 in a disease-relevant neuronal context, we used the preparations of iPSC-derived neurons described above. NPC1 and control neurons were exposed to the BET inhibitor JQ1 (0.3 µM, 1 µM, 3 µM) or vehicle at 30 days after differentiation, and inspected at 24 and 72 hours (Fig. 3; Suppl Fig 1). No substantial differences in neuronal morphology or viability were observed between control and NPC1 neurons at these time points, indicating that JQ1 treatment was well tolerated under these conditions.

**Figure 3:**
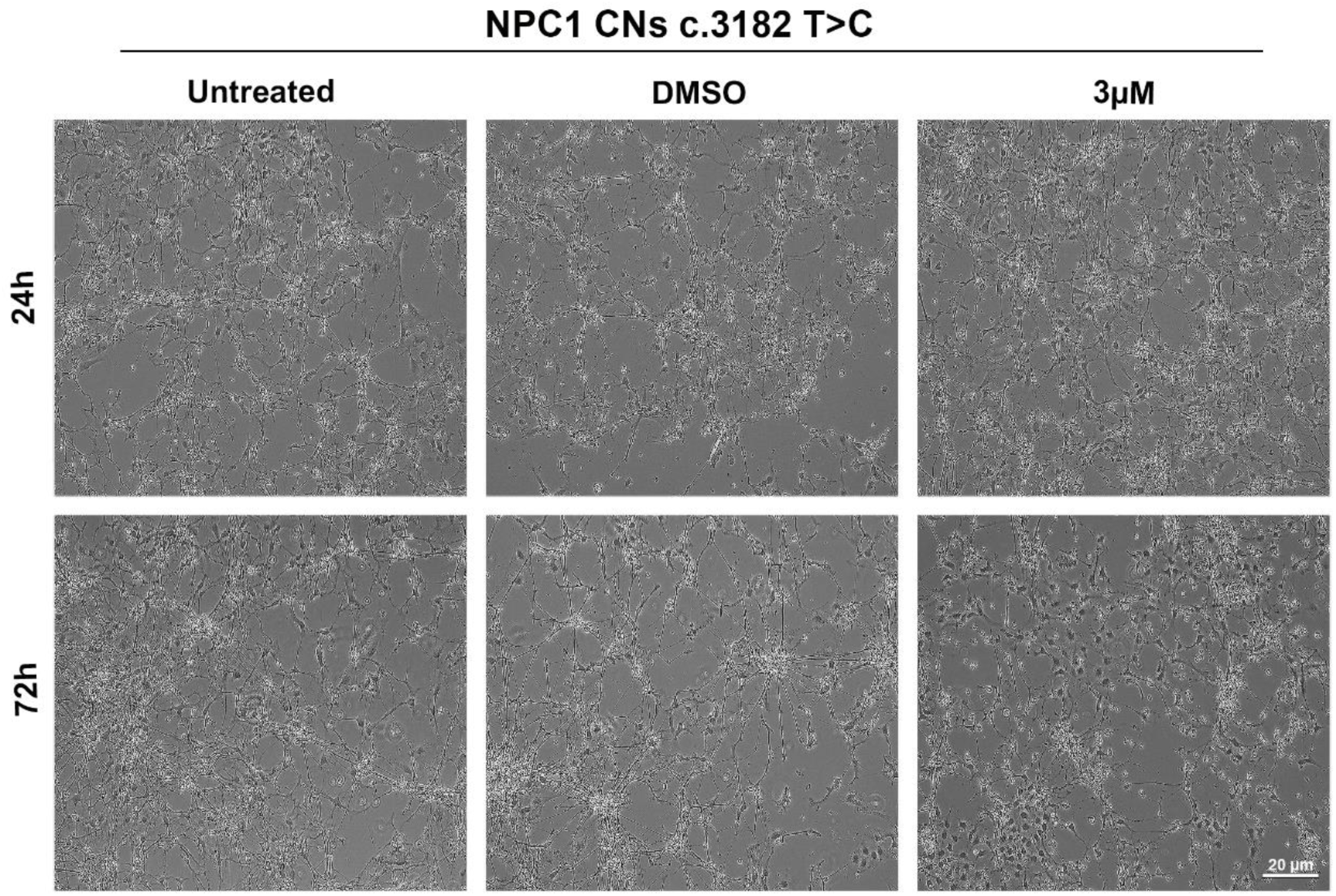
Normal morphology of NPC1 and control iPSC-derived cortical neurons following treatment with the BET inhibitor JQ1. Representative phase-contrast micrographs of iPSC-derived NPCD neurons cultured under indicated experimental conditions (untreated, DMSO used as vehicle and 3 µM JQ1 treatment). Scale bar: 20 μm.

### 4. JQ1 reduces Cholesterol Accumulation in NPC1 Neurons

Next, we assessed whether JQ1 treatment affected intracellular accumulation of non-esterified cholesterol, a cellular hallmark of NPCD, using cytochemical staining with filipin. NPC1 neurons exhibited a marked increase in filipin-positive signal, in accordance with impaired cholesterol trafficking. JQ1 treatment reduced non-esterified cholesterol levels in a time-dependent manner, with the most pronounced reduction observed in NPC1-deficient cortical neurons at later time points (72h) (Fig. 4A, Suppl Fig 2). Co-staining with the neuronal marker βIII-tubulin confirmed that filipin-positive signal was localized within neurites and cell bodies of morphologically intact NPC1 neurons at both 24h and 72h (Fig. 4B, Suppl Fig 2). Quantitative analysis of filipin fluorescence intensity confirmed increased filipin signal in untreated NPC1 neurons compared to control cells (Fig. 4C) and its dose- and time-dependent reduction in NPC1 neurons (Fig. 4D).

**Figure 4:**
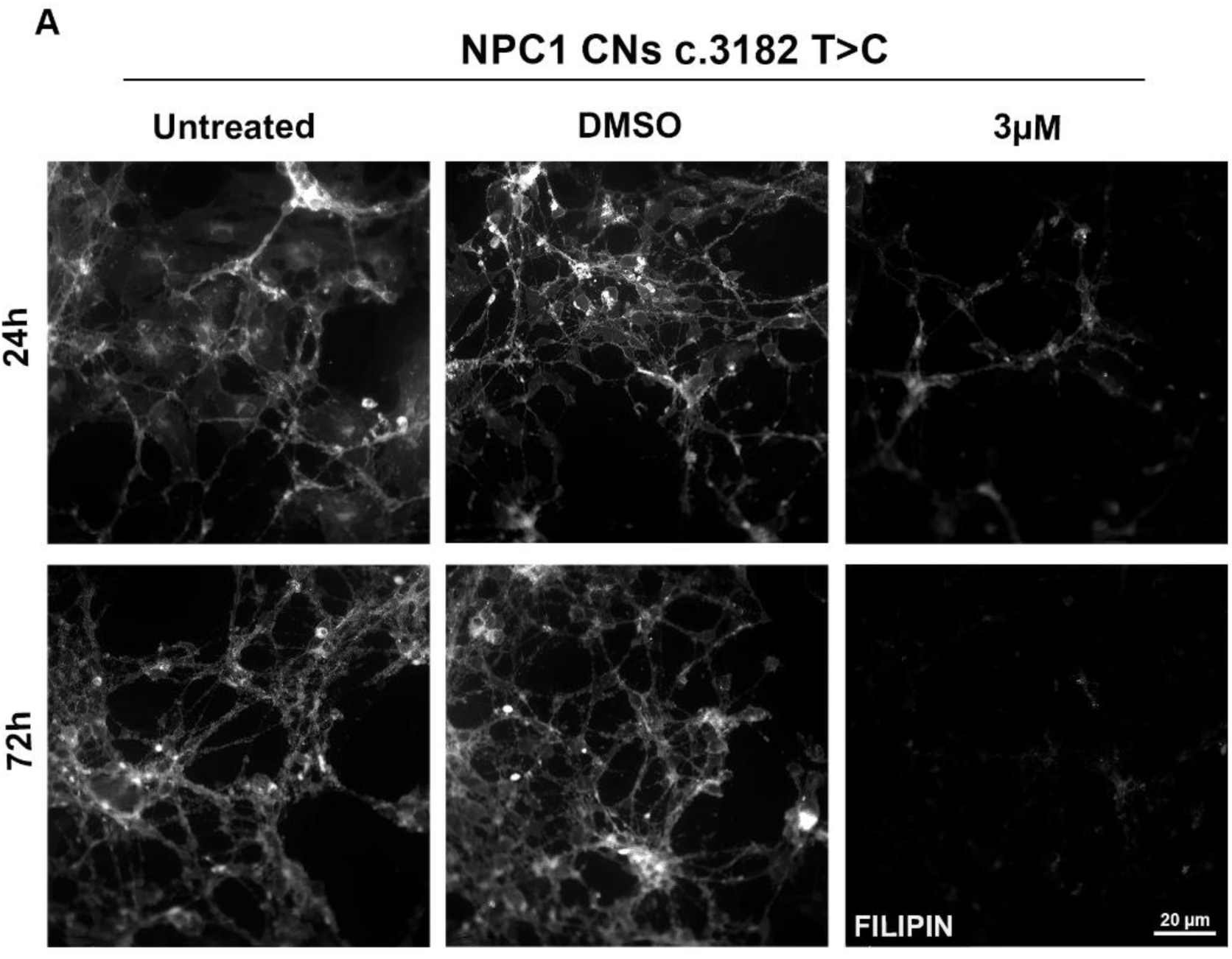
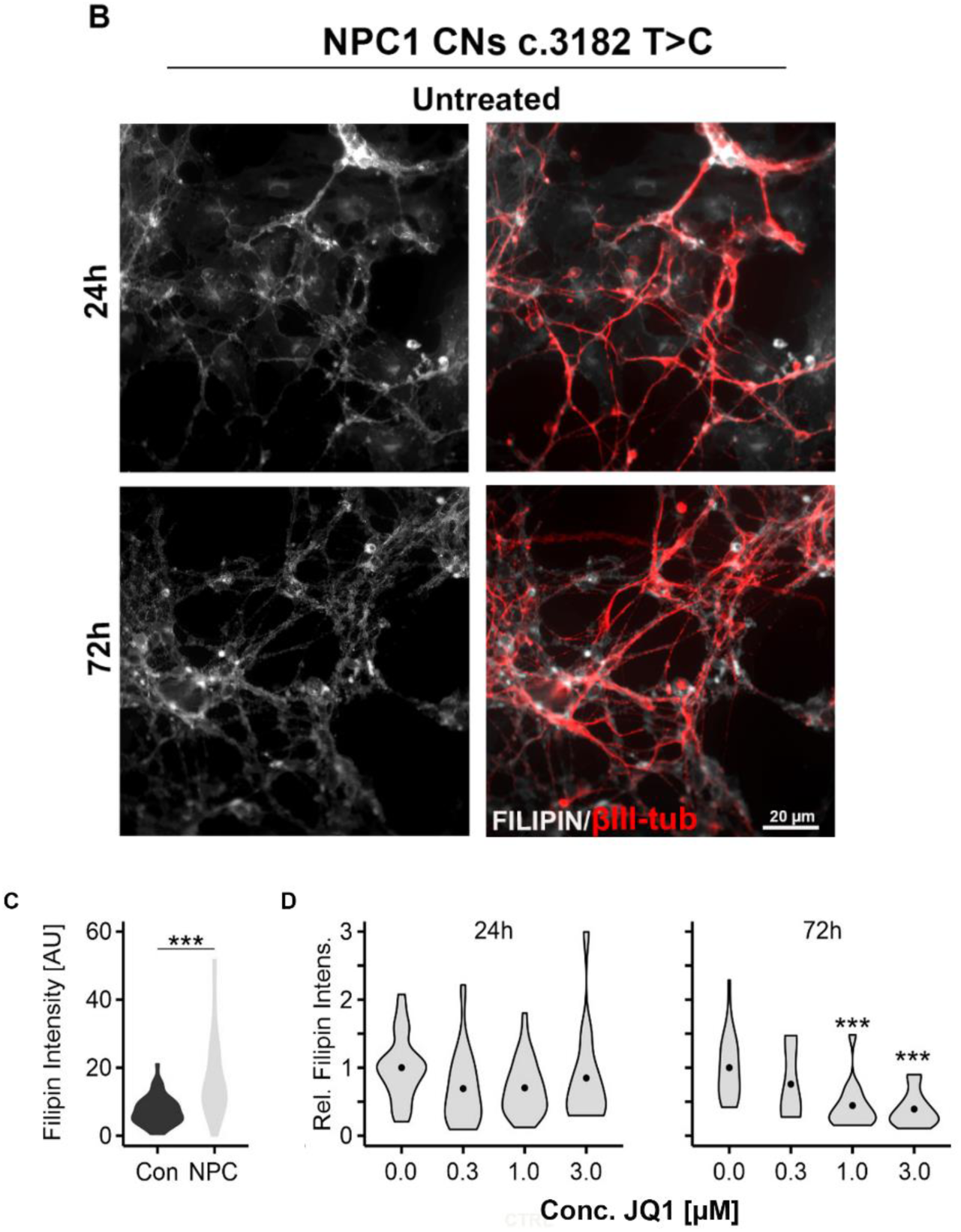
JQ1 induced reduction of cholesterol accumulation in NPC1 iPSC-derived neurons. **(A)** Representative confocal fluorescence micrographs of NPC1 cortical neurons cultured under indicated conditions (untreated, DMSO used as vehicle and 3 µM JQ1 treatment) following immunocytochemical and cytochemical staining for Filipin (grayscale). **(B)** Representative confocal fluorescence micrographs of NPC1 cortical neurons showing Filipin (grayscale) and merged with βIII-tubulin (red) in untreated NPC1 neurons at 24 h and 72 h, highlighting neuronal morphology and cholesterol distribution across time points. Scale bar: 20 μm. **(C-D)** Violin plots showing intensity of filipin fluorescence in untreated control and NPC1 neurons (X) and relative intensities normalized to vehicle (DMSO) cultures after treatment with vehicle (DMSO) or JQ1 at indicated concentrations and durations (Y). Asterisks indicate statistically significant changes compared to vehicle (*** p < 0.001; one-way ANOVA with Tukey’s post hoc test).

### 5. JQ1 Treatment Reduces Cell Death of NPC1 Neurons

TUNEL assays were performed to evaluate neuronal viability in NPC1 iPSC-derived neurons at 24 and 72h of treatment with JQ1 (Fig. 5A, B, Suppl Fig 3). Untreated NPC1 neurons showed a nearly two-fold higher percentage of apoptotic cell death compared to untreated neurons. JQ1 treatment reduced the number of TUNEL-positive cells, notably NPC1 neurons, in a time- and dose-dependent manner (0.3 µM, 1 µM, 3 µM; data not shown), with the strongest effect observed at 72 h (Fig. 5 C,D, Suppl Fig 3).

**Figure 5:**
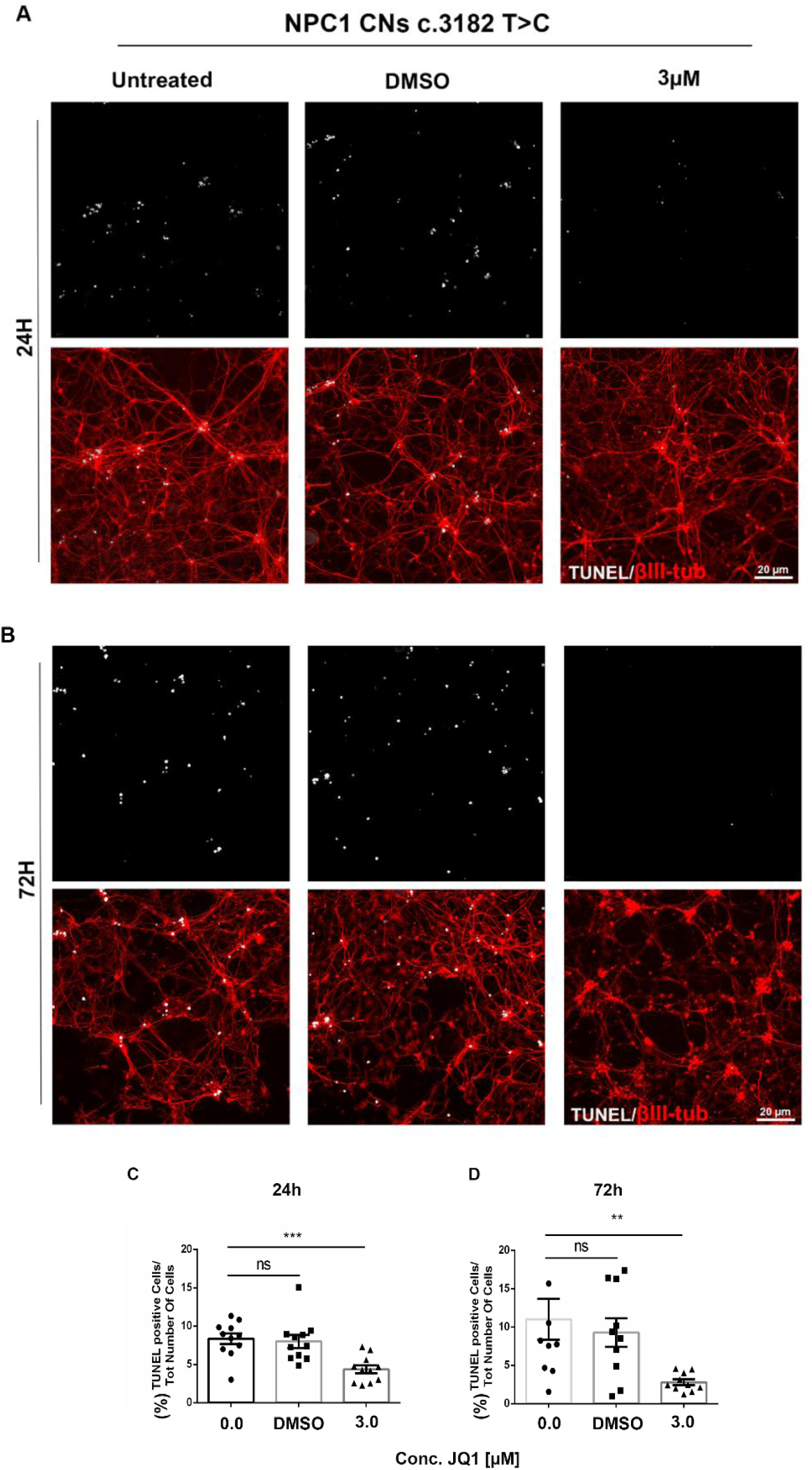
Reduction of cell death in NPC1 iPSC-derived neurons by JQ1 treatment. (A,B) Representative confocal fluorescence micrographs of NPC1 cortical neurons cultured under indicated conditions followed by immunocytochemical staining of βIII-tubulin (red), TUNEL staining (grayscale) at 24h and 72h of treatment. Scale bar: 20 µm. **(C, D)** Column plots (mean ± SEM) showing the percentage of TUNEL-positive NPC1 neurons after 30 days of differentiation and treatment with vehicle (DMSO) and JQ1, at the indicated concentrations and times. (**p ≤ 0.01, ***p ≤ 0.001 vs untreated patient; n = 3; one-way ANOVA with Tukey’s post hoc test).

### 6. JQ1 Treatment Enhance NPC1 Levels in Neurons

To assess the molecular effects of JQ1, the levels of NPC1 were analysed in iPSC-derived neurons by western blotting. Previous studies showed an JQ1-induced increase in cell lines [91], fibroblasts and *NPC1* I1061T mutant mice [92]. Similar as in other models, JQ1 treatment increased NPC1 protein expression in a time- and dose-dependent manner, with the strongest effect observed at the highest dose and the longest treatment (Fig. 6).

**Figure 6:**
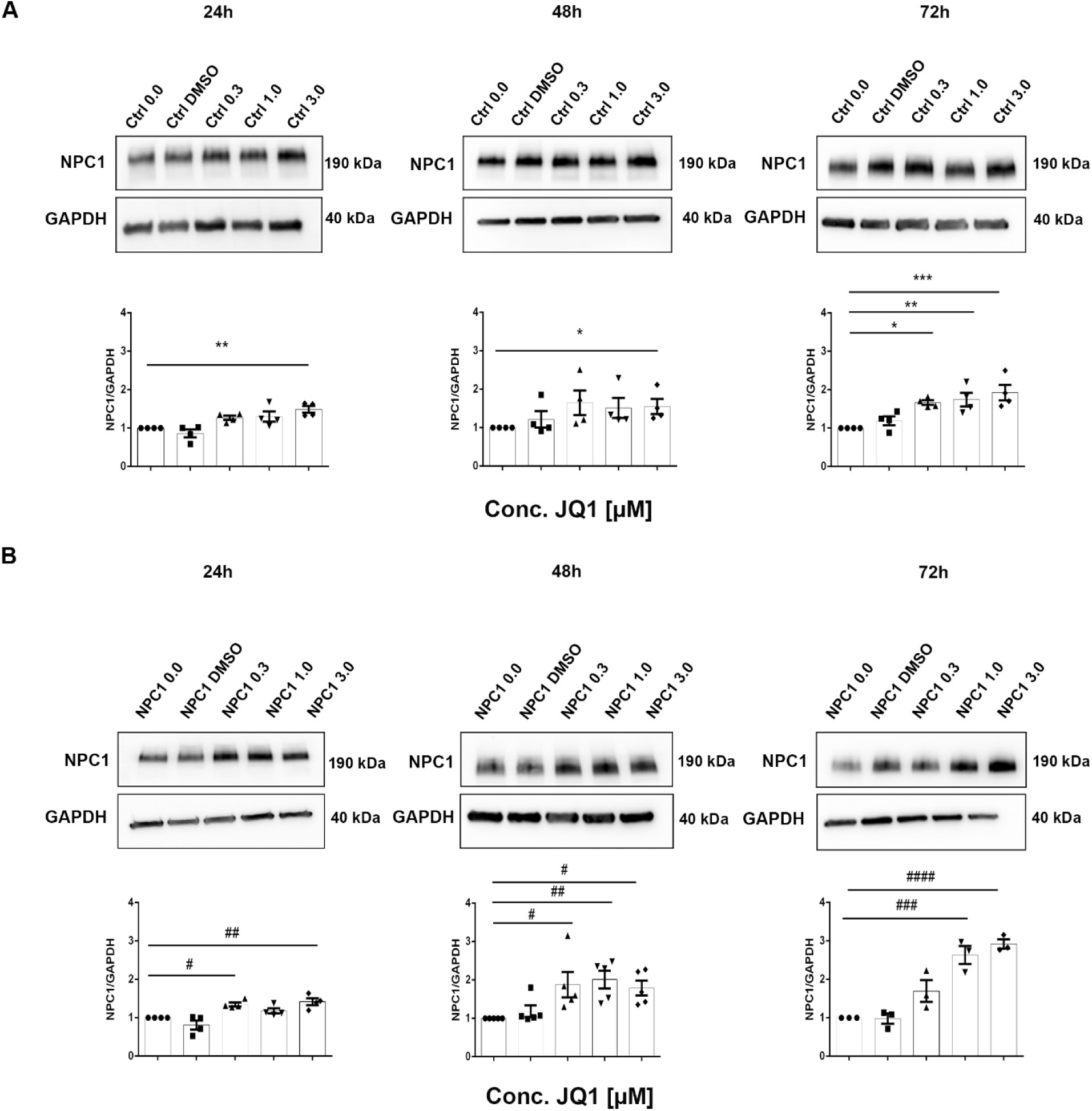
Time- and dose-dependent increase in neuronal NPC1 protein levels after treatment with JQ1. Western blot images and column plots (mean ± SEM) showing levels of NPC1 (190 kDa) and GAPDH (40 kDa) protein in control **(A)** and NPC1 neurons **(B)** after treatment at indicated doses and time periods with JQ1. Levels were normalized to loading control and then to mean levels of untreated cells. Symbols indicate statistically significant changes compared to untreated controls (*p ≤ 0.05, **p ≤ 0.01, ***p ≤ 0.001 vs untreated control; #p ≤ 0.05 ##p ≤ 0.01, ###p ≤ 0.001, ####p ≤ 0.0001 vs untreated patient. n = 3; one-way ANOVA with Tukey’s post hoc test).

## Material and methods

### Derivation and maintenance of iPSCs

The NPCD iPSC line (GM28227), which was derived from a patient fibroblast line (GM18453), and a line from a health donor (AG28851) were purchased from the Coriell Biobank. iPSCs were cultured on 6-well plates coated with Geltrex (A1413202, Gibco) and maintained in mTeSR1 Plus medium (100-0276, Stem Cell Technologies) at 37 °C in a humidified atmosphere containing 5% CO₂. The culture medium was replaced every two day, and cells were passaged upon reaching 70–80% confluence before being differentiated into cortical neurons.

### In Vitro Trilineage Differentiation Assay

Trilineage differentiation was performed using the STEMdiff Trilineage Differentiation Kit (05230, STEMCELL Technologies). According to the protocol, iPSCs were plated on Geltrex with 10 μM Y 27632 (HY-10583, MedChem) and then treated with specific endoderm and mesoderm differentiation media for 5 days, or with ectoderm differentiation media for 7 days.

### Differentiation of iPSCs into cortical neurons

The neuronal differentiation protocol was adapted from a previous publication [98]. Briefly, iPSCs were plated in 6-well plates pre-coated with Geltrex. When cells reached a confluence of 30%, iPSCs medium was replaced with NeuroCult NS-A Basal Medium Human (05750, Stem Cell Technologies) for 16 days. On day 17 of the differentiation period, the medium was replaced with Neurobasal Medium plus (A35829-01, Gibco), supplemented with N-2 (17502001, Gibco) and B-27 Plus supplements (A35828-01, Gibco), 10 ng/mL recombinant human brain-derived neurotrophic factor (BDNF, 450–02, PeproTech), 10 ng/mL recombinant human glial-derived neurotrophic factor (GDNF, 450–10, PeproTech) and 10 ng/mL recombinant human insulin growth factor -1 (IGF1, 10011, PeproTech) until day 24. From day 25 to 30, Neurobasal medium was replaced with BrainPhys Neuronal Medium (05790, Stem Cell Technologies) with the addition of the following factors: NeuroCult SM1 neuronal supplement (05711, Stem Cell Technologies), N-2 supplement, 200 μM ascorbic acid (A4403, Sigma Aldrich), 20 ng/mL recombinant human glial derived neurotrophic factor (GDNF, 450–10, PeproTech), 20 ng/mL recombinant human brain derived neurotrophic factor (BDNF, 450–02, PeproTech). During differentiation, cells were cultured at 37°C, 5% CO2 and 21% O2.

### Immunohistochemical staining

Cells were fixed with 4% paraformaldehyde (PFA) in PBS for 10 min at room temperature (RT), washed with PBS and treated with the blocking and permeabilizing solution containing 5% BSA, and 0.1% Triton X-100 in PBS for 1 h, at RT. Primary antibodies used include: anti-SOX17 (1:3200, O/N at 4°C, rabbit, 81,778, Cell Signaling), anti-Brachyury (1:1600, O/N at 4°C, rabbit, 81,694, Cell Signaling), anti-NCAM (1:400, O/N at 4°C, rabbit, 89,861, Cell Signaling), 1:400 anti-OCT4 monoclonal antibody (MA5-14845, Thermo Fisher Scientific), 1:200 anti-SOX2 monoclonal antibody (149811-82, Thermo Fisher Scientific), 1:250 anti-SSEA4 monoclonal antibody (MA1-021, Thermo Fisher Scientific), 1:100 anti-TRA-1-60 (SC21705, Santa Cruz Biotechnology), 1:200 anti-TBR1 (31940, Abcam), and the neuronal marker 1:500 anti-β III-tubulin monoclonal antibody (T2200, Merck, Germany). Secondary antibodies were conjugated with anti-mouse or anti-rabbit Alexa fluor 488 or 555 (A11070, A21425, Thermo Fisher Scientific) and nuclei were counterstained with 1 µg/ml Hoechst (33342, Thermo Fisher Scientific) at 1:10.000 in PBS for 10 min at RT. Coverslips were mounted using 1:1 PBS-Glycerol.

### Alkaline phosphatase assay

Alkaline phosphatase (ALP) staining was performed following the manufacturer’s instructions (86R 1, Merck KGaA). Briefly, cells were incubated at RT for 30 min with a solution containing naphthol AS-BI and fast red violet LB (86R-1KT, Sigma Aldrich). The cells were photographed using a Leica DM1000 (Leica Microsystems, Wetzlar, Germany) equipped with Leica LAS X software (Leica Microsystems).

### Drug treatment

Cortical neurons at day 30 of differentiation were treated with vehicle (DMSO, final concentration 0.1% v/v [14.1 mM]; D8418, Sigma-Aldrich) or the BET inhibitor (+)-JQ1 (SML1524, Sigma-Aldrich/Merck) at final concentrations of 0.3, 1, and 3 μM for up to 72 h, after dilution from respective stock solutions (1 mM and 3 mM in DMSO). The culture medium was replaced daily with fresh medium containing the same concentrations of JQ1. Control conditions included untreated cells and vehicle-treated cells (DMSO). Cells were collected at 24, 48, and 72 h post-treatment for subsequent analyses. At each time point, cell lysates were harvested and cells grown on glass slides were subjected to immunocytochemical staining.

### TUNEL assay

For TUNEL assays (G3250, Promega) cultured on glass slides were fixed with 4% formaldehyde in PBS for 10 min at RT. Cells were permeabilized by 0.1% Triton X-100 for 10 min at 4 °C, then washed with PBS. equilibration buffer was added to samples for 5 min at RT, prior to incubation with 45 µl equilibration buffer, containing 5 µl Nucleotide Mix and 1 µl TdT, for 1 h at 37 °C. The reaction was stopped by adding 2X SSC for 15 min and nuclei were contrasted by Hoechst (1: 10.000 for 10 min). The percentage of apoptotic and the total cells’ number was evaluated and GraphPad Prism was used to calculate the mean values ± SEM.

### Filipin assay

Filipin was used to label non-esterified cholesterol in cells [99]. Neurons at 30 days of differentiation were fixed with 4% paraformaldehyde (PFA) in PBS for 10 min at RT, followed by washing with PBS. Staining was performed under non-permeabilizing conditions using filipin at 50 ng/mL (freshly prepared from a 10 mg/mL stock solution in ethanol and diluted 1:200 in PBS) and incubated for 2 h at room temperature in the dark. After staining, cells were washed with PBS and subjected to immunocytochemical staining using a monoclonal anti-βIII-tubulin primary antibody (T8578, Sigma-Aldrich). Samples were then immediately imaged by fluorescence microscopy.

### Immunoblotting

Cells were lysed in a solution containing RIPA (S-R0278, Sigma) protease inhibitor cocktail (Roche) and 0.5 mM sodium orthovanadate. Cell extracts were separated by 7.5% sodium dodecyl sulfate-polyacrylamide gel electrophoresis and transferred to nitrocellulose membranes (1610181, Bio-Rad). Blots were blocked with 5% non-fat milk powder in PBS and incubated with specific primary antibodies diluted in blocking solution (1:1000 NPC1, NB400-148, NovusBios and 1:1000 GAPDH, sc-32233, Santa Cruz). Following incubation with secondary antibodies (111-035-003, Jackson ImmunoResearch), proteins were detected by SuperSignal West Pico Chemiluminescent Substrate (Pierce Biotechnologies). Image J software was used to perform immunoblotting analysis.

### Light and Confocal Microscopy

Phase-contrast and confocal images were acquired using a widefield DMi8 microscope (Leica Microsystems) and a confocal Fluoview FV3000 (Evident-Olympus) microscope platform equipped with 405 nm-488 nm-561 nm and 640 nm diode lasers, respectively. Image analysis was carried out using the *open-source* ImageJ software. Acquisition settings (i.e., laser power, beam splitters, filter settings, pinhole diameters and scan mode) were the same for all examined samples of each staining. Representative images were captured and assembled using Adobe Photoshop CS6 software (Adobe Systems Inc.).

### Data Analysis and Visualization

Statistical analyses were performed using Prism software (GraphPad Software, Inc.) and custom written R routines followed by parametric (Student’s t test, one-way ANOVA) or non-parametric (Kruskal-Wallis) tests, to compare sample groups. For IF, Filipin staining, TUNEL assay and WB, a minimum of 3 technical and 3 biological replicates were performed for all experiments.

## Discussion

In this study, we established and characterized a human iPSC-derived cortical neuronal model of NPCD carrying the frequent p.I1061T NPC1 variant, and used it to investigate the effects of epigenetic modulation via BET proteins. NPCD patient-derived iPSCs maintained key pluripotency features, including ESC-like morphology, expression of pluripotency markers, and trilineage differentiation capacity, consistent with previous reports [72,74,75,77,95]. Following neural differentiation, both control and NPC1 iPSCs generated cortical neurons expressing TUJ1 and TBR1 [70,76,99]. NPCD-derived neurons exhibited overall comparable morphology and network organization to controls, indicating that the p.I1061T variant does not significantly impair early neuronal differentiation. Despite this, NPC1 neurons displayed a marked accumulation of unesterified cholesterol, as demonstrated by filipin staining, consistent with defective lysosomal cholesterol export [17,62,67,76]. The latter is a central hallmark of NPCD and is associated with downstream lysosomal dysfunction, impaired autophagy, and cellular stress [26,29,30,38,45,48,49]. In line with these alterations, NPC1 neurons showed increased susceptibility to apoptosis, as indicated by TUNEL analysis, consistent with previous studies in NPC1 cellular and iPSC-based models [29,35,40,50,76,99]. In addition, western blot analysis revealed reduced NPC1 protein levels in patient-derived neurons consistent with mutation-associated misfolding, endoplasmic reticulum retention, and enhanced proteasomal degradation [93,94,97]. Together, these phenotypes: cholesterol accumulation, reduced NPC1 protein levels, and increased apoptosis, recapitulate key aspects of NPCD and support the validity of this iPSC-derived neuronal model for mechanistic and therapeutic studies [70,72,74–76,99]. To our knowledge, this is the first study to examine the effects of BET inhibition in a human iPSC-derived neuronal model of NPCD. Pharmacological BET inhibition with JQ1 reduced intracellular cholesterol accumulation and apoptosis while increasing NPC1 protein levels, with maximal effects observed after 72 hours of treatment. Importantly, no overt toxicity or major morphological changes were observed under these conditions. The increase in NPC1 protein levels following BET inhibition is particularly notable. Whether other, NPC1-independent pathways contribute to the positive effects is not known. The pleiotropic nature of BET proteins means that prolonged or systemic inhibition may have broader effects not captured in the present *in vitro* model [83,87,88]. Future studies in more complex systems are required to define mechanism, specificity, and translational relevance. BET proteins, particularly BRD4, are known regulators of transcriptional programs involved in autophagy, lysosomal function, inflammation, stress responses, and lipid metabolism [81,87–91]. Although BET dysregulation has not previously been described in NPCD, our findings suggest a potential role for BET-dependent regulation in neuronal lipid homeostasis. The reduction in cholesterol accumulation following JQ1 treatment further supports the idea that BET inhibition may influence intracellular cholesterol handling, potentially via modulation of lysosomal and autophagic pathways [89,91,92]. However, the precise mechanisms underlying these effects, whether involving improved lysosomal function, altered cholesterol trafficking, or enhanced proteostasis, remain to be determined.

Overall, our findings suggest that BET inhibition targets multiple pathways involved in NPCD pathology, supporting epigenetic regulation as an upstream mechanism of disease. Given the limited efficacy of current therapies in preventing disease progression [58–65], BET inhibition may represent a promising complementary therapeutic strategy, pending further validation in additional experimental models [78,87–92]. In conclusion, this study demonstrates that patient-derived iPSC cortical neurons provide a relevant human model for NPCD and supports the hypothesis that epigenetic modulation via BET inhibition can ameliorate key disease-associated phenotypes, including cholesterol accumulation, reduced NPC1 protein levels, and increased apoptosis. These findings provide a rationale for further investigation of BET proteins as potential modulators of NPC1-associated neuronal dysfunction.

## Ethic Statement

The biological material from the subjects included in this study was collected following procedures in accordance with the ethical standards of the declaration of Helsinki protocols.

## Authors’ contribution

F.B., C.C, F.W.P. and V.P. drafted the manuscript. F.B., M.P., N.C. and A.B. performed the experiments and analyzed the data. C.C., R.B., C.M., S.P., M.T., F.W.P. and V.P. analyzed the data and critically revised the manuscript. All authors have read and agreed to the published version of the manuscript.

