## Supplemental Figures for "Pharmacological BET Protein Inhibition Reverts Niemann-Pick Type C Disease-Associated Changes in Human iPSC-Derived Cortical Neurons"

### Supplemental Figures Benigni et al.

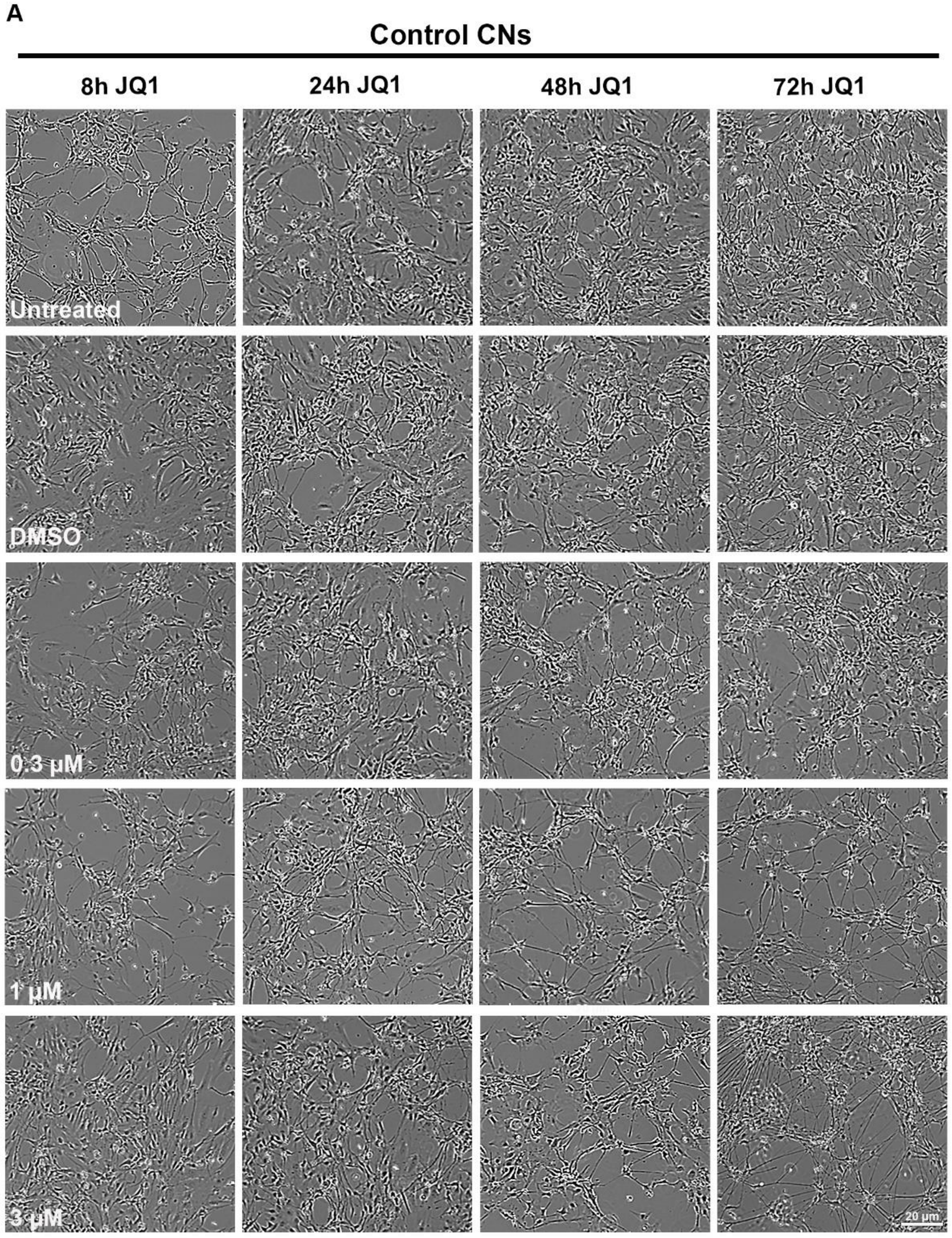

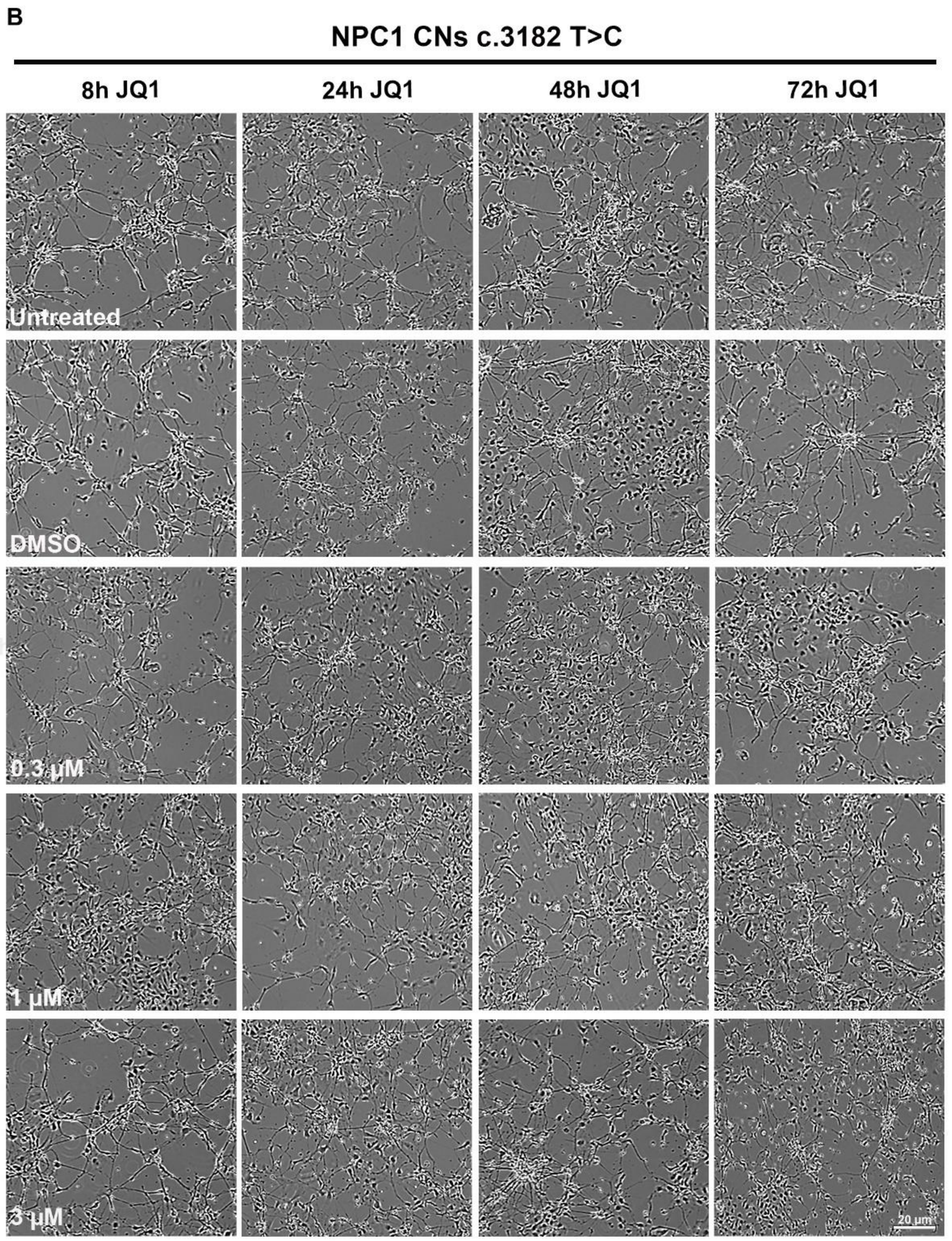


**<u>Supplemental Figure 1</u>. *Representative micrographs of 30-day differentiated iPSC-derived cortical neurons treated with the BET inhibitor JQ1.*** *Rows indicate experimental conditions (untreated, vehicle control DMSO, and JQ1 at 0.3, 1, and 3 µM), while columns indicate treatment duration (8, 24, 48, and 72 hours). In panel A control cortical neurons are shown, while in panel B patient-derived cortical neurons are shown. Scale bar: 20 µm.*

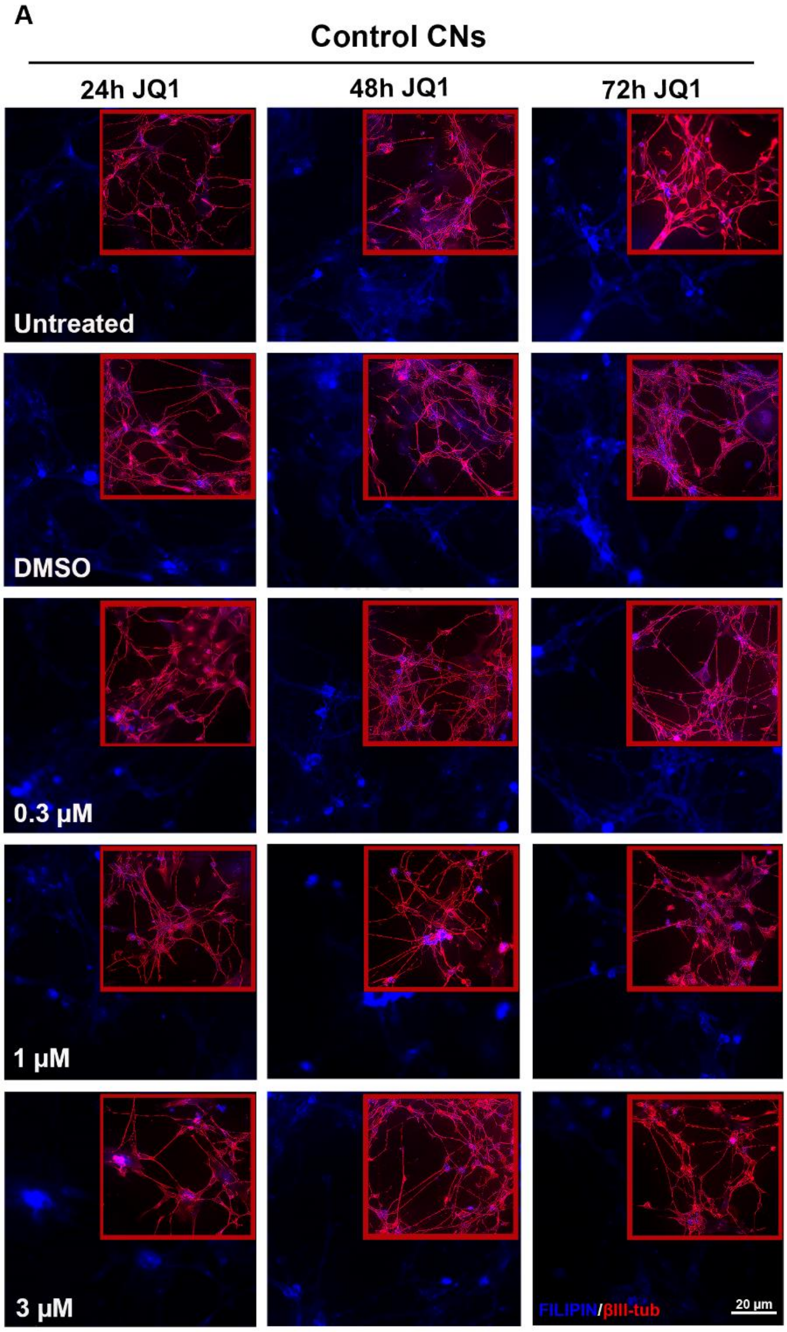
A
Control CNs
24h JQ1
48h JQ1
72h JQ1
Untreated
DMSO
0.3 µM
1 µM
3 µM
FILIPIN/βIII-tub
20 µm

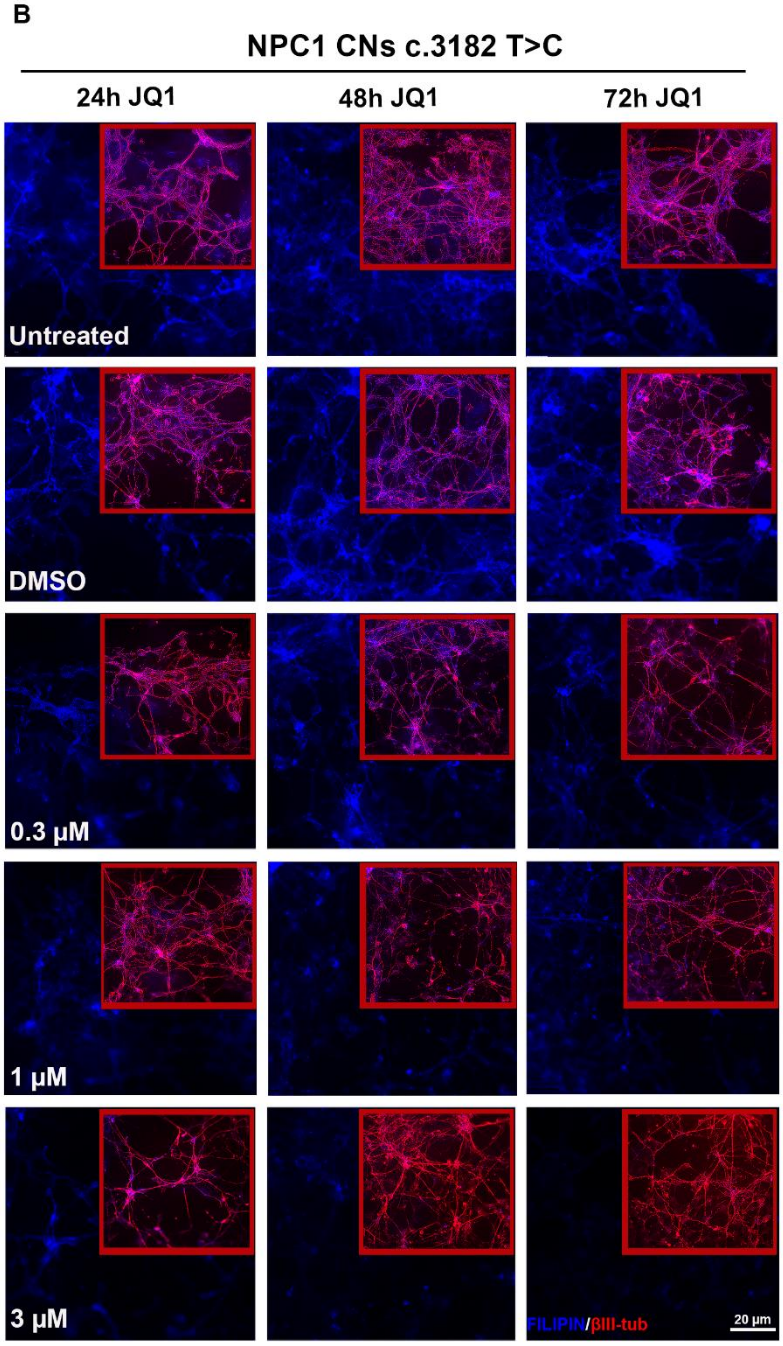


**<u>Supplemental Figure 2</u>. Representative immunofluorescence images of control (panel A) and NPC1 (panel B) cortical neurons for βIII-tubulin (red) and Filipin (blue) after 24h, 48h and 72h JQ1 treatment.** *In the red boxes, representative confocal images of neurons differentiated for 30 days and stained with Filipin (blue) and βIII-tubulin (red) under the indicated conditions: untreated, vehicle control DMSO, and JQ1 (0.3, 1, 3 μM). JQ1 reduced Filipin intensity in a dose-dependent manner, with the strongest effect observed in NPC1 patient neurons. Scale bar: 20 μm.*

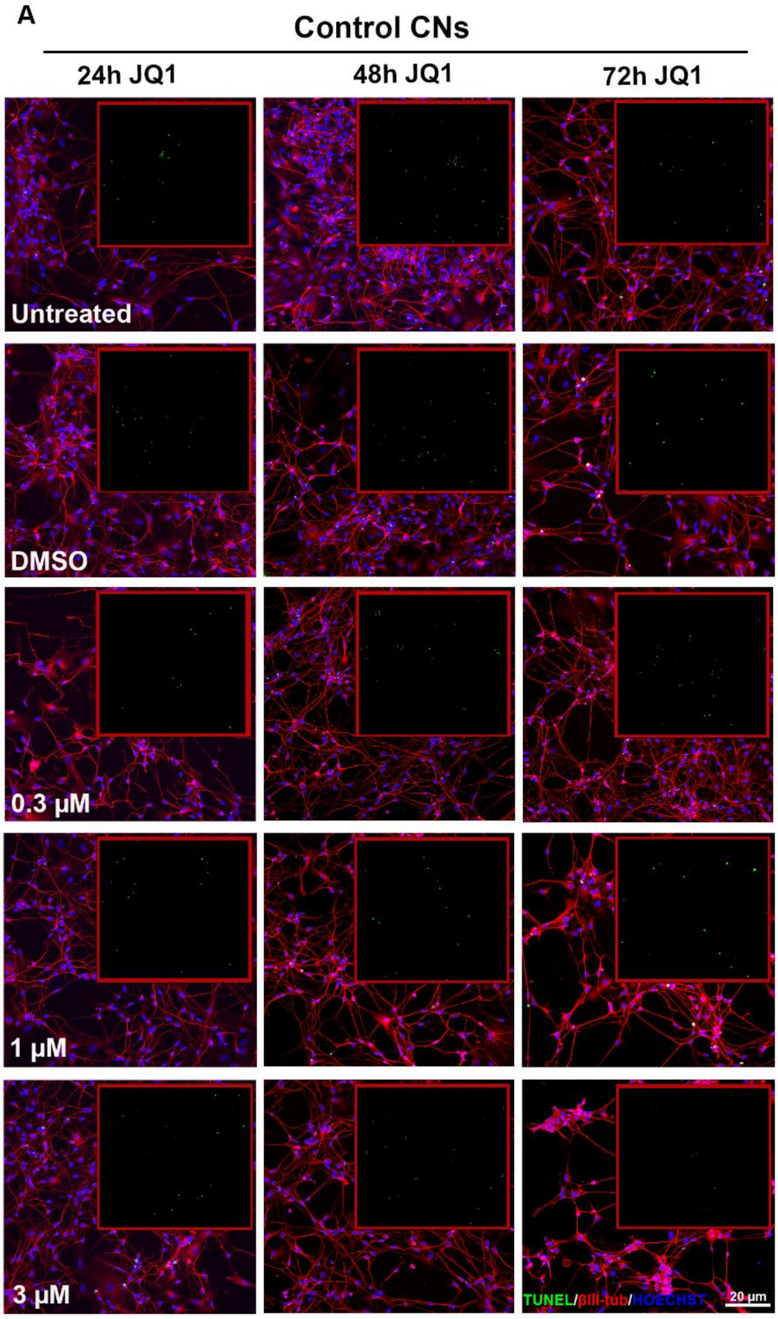
A
Control CNs
24h JQ1
48h JQ1
72h JQ1
Untreated
DMSO
0.3 μM
1 μM
3 μM
TUNEL/βIII-tub/HOECHST
20 μm

B

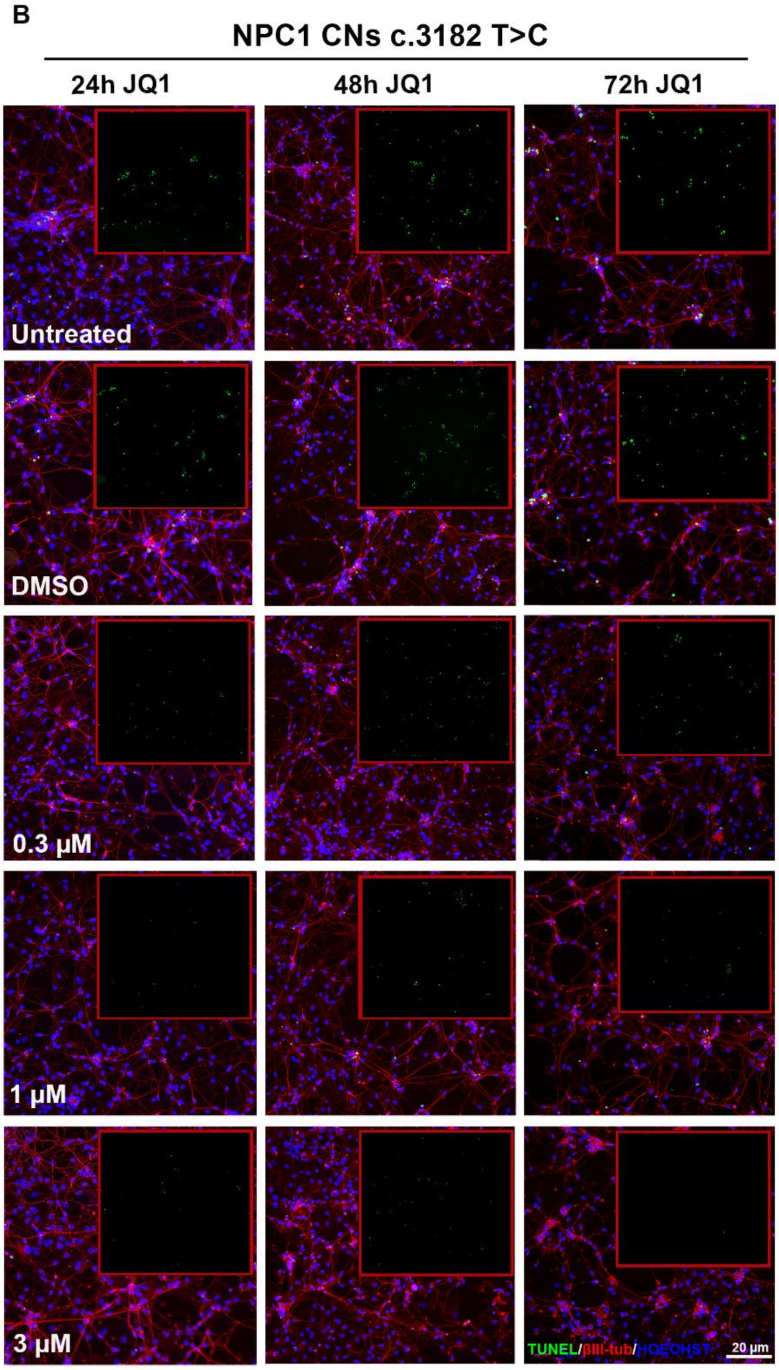
NPC1 CNs c.3182 T>C
24h JQ1
48h JQ1
72h JQ1
Untreated
DMSO
0.3 µM
1 µM
3 µM
TUNEL/βIII-tub/HOECHST
20 µm

C

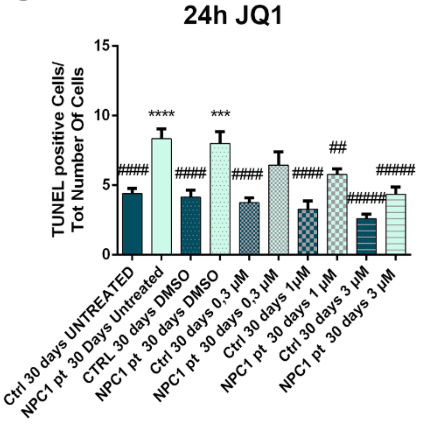
24h JQ1
TUNEL positive Cells/
Tot Number Of Cells
15
10
5
0
Ctrl 30 days UNTREATED
NPC1 pt 30 Days Untreated
CTRL 30 days DMSO
NPC1 pt 30 days DMSO
Ctrl 30 days 0,3 µM
NPC1 pt 30 days 0,3 µM
Ctrl 30 days 1µM
NPC1 pt 30 days 1 µM
Ctrl 30 days 3 µM
NPC1 pt 30 days 3 µM

D

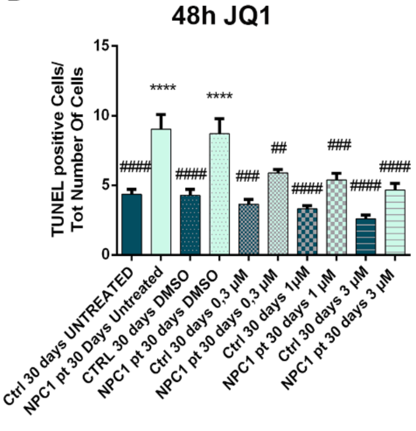
48h JQ1
TUNEL positive Cells/
Tot Number Of Cells
15
10
5
0
Ctrl 30 days UNTREATED
NPC1 pt 30 Days Untreated
CTRL 30 days DMSO
NPC1 pt 30 days DMSO
Ctrl 30 days 0,3 µM
NPC1 pt 30 days 0,3 µM
Ctrl 30 days 1µM
NPC1 pt 30 days 1 µM
Ctrl 30 days 3 µM
NPC1 pt 30 days 3 µM

E

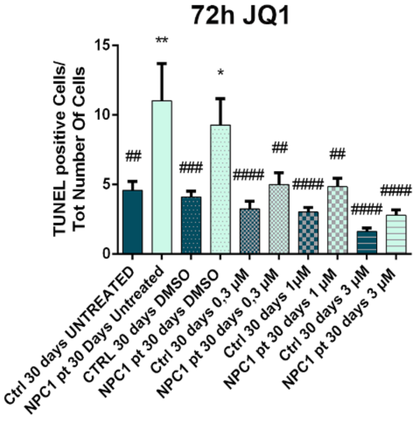
72h JQ1
TUNEL positive Cells/
Tot Number Of Cells
15
10
5
0
Ctrl 30 days UNTREATED
NPC1 pt 30 Days Untreated
CTRL 30 days DMSO
NPC1 pt 30 days DMSO
Ctrl 30 days 0,3 µM
NPC1 pt 30 days 0,3 µM
Ctrl 30 days 1µM
NPC1 pt 30 days 1 µM
Ctrl 30 days 3 µM
NPC1 pt 30 days 3 µM

***<u>Supplemental Figure 3</u>. Representative immunofluorescence images of control (panel A) and NP C1 (panel B) cortical neurons for βIII-tubulin (red), TUNEL (green) and Hoechst (blue) after 24h, 48h and 72h JQ1 treatment.*** *In the red boxes, representative staining for TUNEL (green). Scale bar: 20 µm. (C,D,E) Quantification of TUNEL-positive cells (%of TUNEL positive cells over the total number of cells) in control (dark bars) and NPC1 (light bars) neurons after 30 days of differentiation, respectively at 24h, 48h and 72h JQ1 treatment. Cells were untreated or treated with vehicle (DMSO) or JQ1 (0.3, 1, 3 µM). Data are mean ± SEM (n = 3); one-way ANOVA with Kruskal–Wallis post-test. *$p \leq 0.05$, **$p \leq 0.01$, ***$p \leq 0.001$, ****$p \leq 0.0001$ vs untreated control; ##$p \leq 0.01$, ###$p \leq 0.001$, ####$p \leq 0.0001$, #####$p \leq 0.00001$ vs untreated patient.*
